# Deep learning in human neurons predicts mechanistic subtypes of Parkinson’s

**DOI:** 10.1101/2022.03.10.482156

**Authors:** Karishma D’Sa, James R. Evans, Gurvir S. Virdi, Giulia Vecchi, Alex Adam, Ottavia Bertolli, James Fleming, Hojong Chang, Dilan Athauda, Minee L. Choi, Sonia Gandhi

## Abstract

Parkinson’s disease (PD) is a common, devastating, and incurable neurodegenerative disorder. Several molecular mechanisms have been proposed to drive PD, with genetic and pathological evidence pointing towards aberrant protein homeostasis and mitochondrial dysfunction. PD is clinically highly heterogeneous, it is likely that different mechanisms underlie the pathology in different individuals, each requiring a specific targeted treatment. Recent advances in stem cell technology and fluorescent live-cell imaging have enabled the generation of patient-derived neurons with different mechanistic subtypes of PD. Here, we performed multi-dimensional fluorescent labelling of organelles in iPSC-derived neurons, in healthy control cells, and in four different disease subclasses. We generated a machine learning-based model that can simultaneously predict the presence of disease, and its primary mechanistic subtype. We independently trained a series of classifiers using both quantitative single-cell fluorescence variables and images to build deep neural networks. Quantitative cellular profile-based classifiers achieve an accuracy of 82%, whilst image based deep neural networks predict control, and four distinct disease subtypes with an accuracy of 95%. The classifiers achieve their accuracy across all subtypes primarily utilizing the organellar features of the mitochondria, with additional contribution of the lysosomes, confirming their biological importance in PD. Taken together, we show that machine learning approaches applied to patient-derived cells are able to predict disease subtypes, demonstrating that this approach may be used to guide personalized treatment approaches in the future.

## INTRODUCTION

Parkinson’s disease (PD) is a progressive neurodegenerative disorder that encompasses several pathogenic processes that converge on the accumulation of misfolded α-synuclein in Lewy bodies and neurites, and degeneration of dopaminergic neurons in the substantia nigra, resulting in an array of motor, cognitive, neuropsychiatric and autonomic deficits (Braak et al., 2003; Spillantini et al., 1997; Weinreb et al., 1996). The age of onset, rate of disease progression and severity of motor and non-motor symptoms display considerable individual variation (reviewed here (Cheng et al., 2010)). This is most likely due to differences in the underlying molecular mechanisms occurring in different subtypes of the disease (reviewed here (Kusumoto and Yuasa, 2019)). Critically, there are currently no approaches to define the molecular heterogeneity, and therefore no opportunity to understand the mechanisms that may drive the different phenotypic subtypes. Furthermore, the lack of mechanistic understanding in PD, and of how best to target the condition, has led to the absence of any cure, or even any disease-modifying therapy to date.

Clinicians have increasingly adopted a patient centric approach to the symptomatic treatment of PD, using multiple drugs and delivery systems according to specific individual needs of patients. However, when considering how to achieve disease modification, the majority of studies still utilize a “one-size-fits-all” approach. This may be fundamentally flawed as it is unlikely that there is a single solution or treatment that works for all patients, and a more “precise” approach based on the molecular underpinnings of the disease in each patient may be the best way forward. Thus, an unmet challenge is to make an early and accurate molecular level diagnosis of the condition, as this would enable the field to consider targeted interventions appropriate to an individual’s condition, and offer an opportunity to do this at the earliest possible time.

To address this challenge, we applied a deep learning approach to cellular models of PD to generate a predictive model of different mechanisms of disease. PD is known to be caused by a complex interplay of genetic and environmental drivers. Two common and critical pathways that drive pathology include (i) the accumulation of insoluble aggregates of the protein alpha-synuclein, implying that protein misfolding and impaired protein homeostasis causes a ‘proteinopathy’, or a ‘synucleinopathy’ (Campbell et al., 2020; Spillantini et al., 1997) and (ii) the accumulation of abnormal mitochondria with impaired bioenergetic function, and reduced mitochondrial clearance (Angelova et al., 2018) and reviewed here (Deas et al., 2011)). To model PD, we utilized patient-derived iPSC derived cortical neurons: these are a robust preclinical cell model for disease, recapitulating the human genomic and proteomic environment of a differentiated cell type that is affected in disease (Laperle et al., 2020; Rowe and Daley, 2019; Takahashi and Yamanaka, 2006). We defined four cellular subtypes that map to both of these two key pathways that lead to disease: (i) patients with mutations in the SNCA gene develop an autosomal dominant aggressive form of Parkinson’s with predominant protein aggregation, that is caused directly by the mutation in the SNCA gene: therefore, iPSC derived neurons from a patient with SNCA triplication were used to model ‘familial proteinopathy’. (ii) protein aggregates are known to spread from cell to cell in the brain in a prion-like manner, inducing proteotoxic stress: we exposed healthy iPSC derived neurons to aggregates of alpha-synuclein to recapitulate the ‘environmental proteinopathy’ (Cremades et al., 2012; Ludtmann et al., 2018). (iii) exposure to pesticides with subsequent mitochondrial complex 1 impairment can induce PD, and patients with PD exhibit widespread impairment of complex 1 dependent respiration: we applied a complex 1 inhibitor, rotenone to generate a model of ‘toxin induced mitochondrial dysfunction’ (iv) mutations in the PINK1 and PARKIN genes cause autosomal recessive early onset PD, and these mutations directly result in impaired clearance of damaged mitochondria (mitophagy), resulting in the accumulation of abnormal mitochondria in neurons (Deas et al., 2011): we applied an inducer of mitophagy, oligomycin/antimycin, to generate a model of ‘Mitophagy induced PD’.

We simultaneously fluorescently labelled specific cellular compartments (the nucleus, mitochondria and lysosomes), and we performed high content live single cell imaging of iPSC derived neurons. Notably, the multiplexed imaging was designed to capture specific organellar dysfunction that has been previously validated for relevance in disease. Using data from 8 different plates (total 1,560,315 cells), we generated a number of models to predict disease state and disease subtype. We generated two broad types of classifier (i) a prediction classifier based on features automatically extracted (56 features), providing deep profiling of cellular phenotypes: this classifier has the advantage of high explainability using the ranking of features (ii) prediction classifiers based on images and deep neural network analyses, utilizing the power of computer vision to extract large amounts of unbiased information: this classifier has very high accuracy, but with less explainability. In this study we compared the two approaches to define the presence of disease within a group of neurons and the origin, or cause, of that disease subtype. Our work identifies the specific features in neurons that are able to predict different cellular subtypes of the disease, and therefore provides valuable biological insights into the mechanisms of PD.

## RESULTS

### Induced pluripotent stem cell derived neurons as a model for human Parkinson’s

Human iPSCs were generated through reprogramming skin cells from healthy donors or PD patients carrying SNCA triplication, which causes a familial early onset aggressive form of PD, with widespread proteinopathy. Neuronal differentiation was performed through dual SMAD inhibition using SB431542 (10 μM, Tocris) and dorsomorphin dihydrochloride (1 μM, Tocris) using a protocol adapted from (Shi et al., 2012), see the schematic in Figure 1a). 20,000 - 40,000 cells were plated onto a 96-well plate at day 50, and maintained until use (> day 60). Upon terminal differentiation, the culture is highly enriched with populations of neurons displaying neuronal markers (MAP2: 90.01 ± 3.516%, Figure 1bi) with both lower-(TRB1: 46.67 ± 4.985%, CTIP2: 47.30 ± 5.037, Figure 1bii) and upper-layer neuronal markers (SATB2: 41.15 ± 4.501%, Figure 1biii). Neuronal cultures responded to physiological concentration of glutamate (5 uM) which induces the opening of potential-sensitive Ca^2+^ channels that are specific for neurons, thus confirming the majority of the population are glutamatergic neurons (62.92 ± 3.819%, Figure 1c). A key hallmark of PD is the accumulation of intraneuronal aggregates, consisting of phosphorylated alpha-synuclein. hiPSC-derived neurons carrying 3x SNCA mutation express more phosphorylated forms of α-Syn compared to control iPSC-derived neurons (Figure 1di & ii).

**Figure 1.**
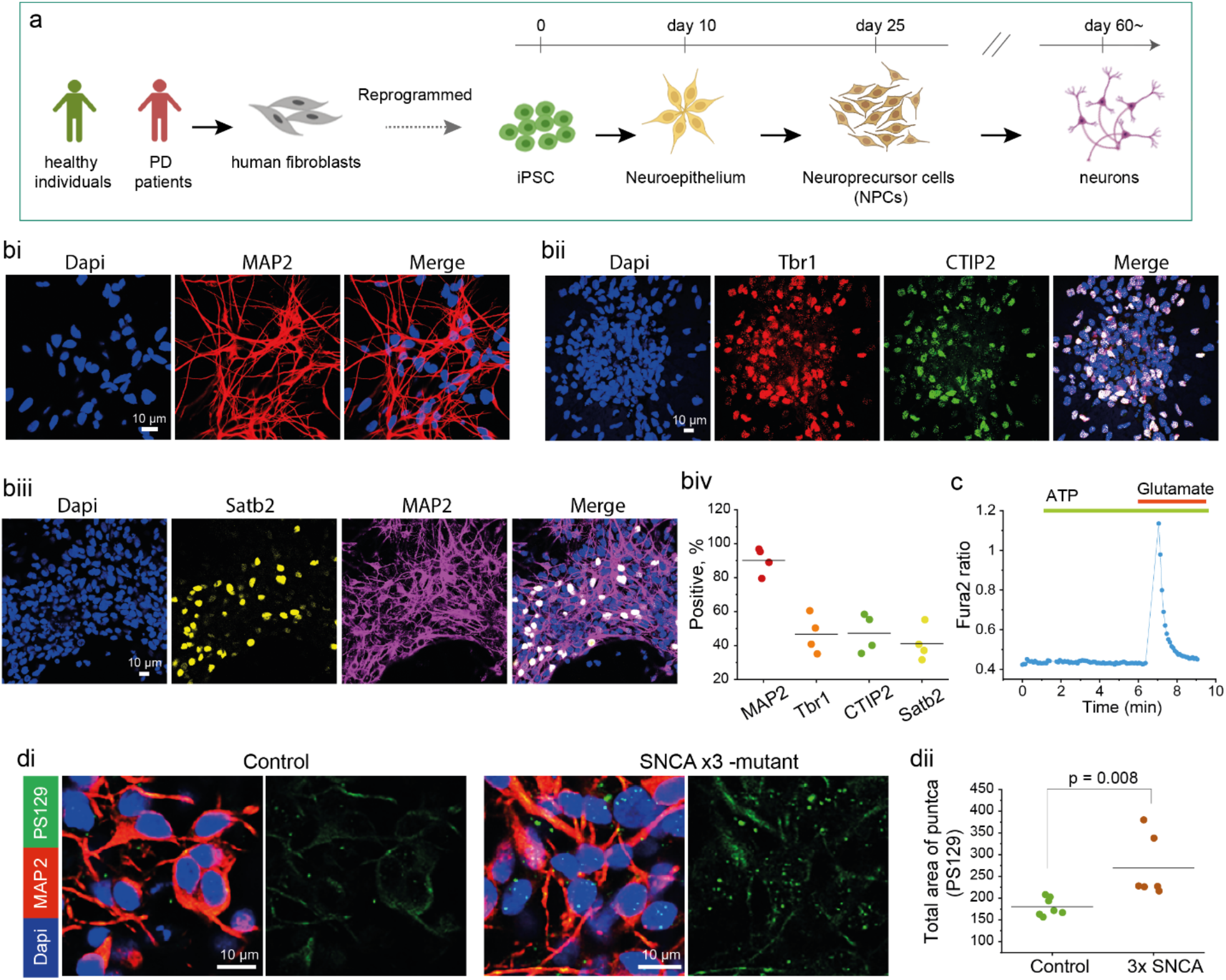
Generation a human PD model using hiPSCs. **a**. Schematic showing hiPSC derived neuronal differentiation strategy. Fibroblasts from patients with PD or healthy donors are reprogrammed into hiPSC and differentiated into cortical neurons using protocol from (Shi et al., 2012). **bi-iv**. Characterization of iPSC derived neurons using ICC for the representative images of i) MAP2, neuronal marker, ii) Tbr1 and CTIP2, deep cortical layer, iii) Satb2, upper cortical layer and iv) the quantification **c**. Calcium imaging measured with Fura-2 shows the hiPSC-derived cortical neurons exhibit calcium signal in response to physiological concentration (5 µM) of glutamate. **di-ii**. hiPSC-derived neurons from PD patients with 3xSNCA mutation display increase phosphorylated α-Syn (aggregated form of α-Syn).

Our data demonstrate that we are able to generate human neurons with the functional and molecular identity of neurons from the human cortex, and that the Subtype 1: familial proteinopathy, exhibits the pathological hallmark of a proteinopathy (Angelova et al., 2020; Ludtmann et al., 2018; Whiten et al., 2018).

### Development of a classifier to identify disease states in PD

Mitochondrial dysfunction and synucleinopathy are two primary pathologies of PD which are induced by various conditions (Gandhi et al., 2009; Hsieh et al., 2016; Spillantini et al., 1997). We established a set of disease states (four subtypes as described in Table 1) led by two primary pathologies; α-Syn aggregation and mitochondrial dysfunction. The 3x SNCA and oligomer disease states belong to the aggregation pathway, while Complex 1 inhibition and impaired mitophagy are associated with mitochondrial toxicity. To establish Subtype 1: a familial proteinopathy, we differentiated neurons with iPSC derived from PD patients carrying 3x SNCA which are reported to have elevated aggregation levels in neurons (Figure 1di & ii) (Angelova et al., 2020; Ludtmann et al., 2018). We also established Subtype 2; an ‘environmental proteinopathy’ by treating cells with α-Syn oligomers, toxic soluble species of α-Syn which results in oligomer induced toxicity (Angelova et al., 2020; Ludtmann et al., 2018; Whiten et al., 2018). For Subtype 3: the mitochondrial toxic state was induced by disrupting the mitochondrial electron transport chain using Rotenone (5 µM, mitochondrial complex 1 inhibitor) (Abramov et al., 2010; Esteras et al., 2017; Reeve et al., 2015). For Subtype 4: mitophagy, a process of eliminating unhealthy mitochondria was induced by co-treating with Oligomycin A (1 µM, Sigma) and Antimycin (1 µM, Sigma) (Soutar et al., 2018). We adopted live-cell imaging of multiplexed dyes to reveal the organelles of the hiPSC-derived neurons (nucleus, mitochondria and lysosomes). In particular, we selected tracker dyes for organelles that are involved in the four subtypes: mitochondria and lysosomes. We demonstrate that the signal from these fluorescent dyes varies depending on the health of the organelles and that this requires live imaging, as such changes of organelle function and organellar networks are not apparent with fixation (Figure 2a & b). Rotenone, a complex 1 inhibitor of electron transport chain (ETC), depolarizes mitochondrial membrane potential labelled with Tetramethylrhodamine, Methyl Ester (TMRM, *p* = 0.0046, Figure 2ai & ii), resulting in loss of fluorescent intensity of TMRM. Lysosomal activity is also elevated by Chloroquine which is labelled with Lysotracker (*p* = 0.0020, Figure 2bi & bii), resulting in an increase in fluorescence.

**Table 1.** Pathological details of the cellular subtypes of PD. Subtype 1: Cells generated with 3x SNCA mutation represent ‘familial proteinopathy’. **Subtype 2:** ‘Environmental proteinopathy’ was induced by exposing cells to exogenous protein aggregates. **Subtype 3:** Toxin-induced mitochondrial dysfunction was obtained by exposing cells to rotenone, a Complex 1 inhibitor. **Subtype 4:** We induced mitophagy using stimulation with oligomycin/antimycin, to represent our fourth subtype.

| Disease states | Experimental conditions | Primary pathology | Common pathology |
| --- | --- | --- | --- |
| Subtype 1 *'Familial proteinopathy'* | Neurons are differentiated from iPSC derived from PD patients carrying 3x SNCA mutation. | Formation of α-Syn aggregation with beta sheet | Neuronal death |
| Subtype 2 *'Environmental proteinopathy'* | Human recombinant α-Syn oligomer (0.5 μM) treatment | | |
| Subtype 3 *'Complex 1 induced mitochondrial dysfunction'* | Rotenone (5 μM) treatment to inhibit Complex 1 | Accumulation of abnormal mitochondria | |
| Subtype 4 *'Mitophagy induced mitochondrial dysfunction'* | Co-treatment with oligomycin (1 μM) and antimycin (1 μM) | | |

**Figure 2.**
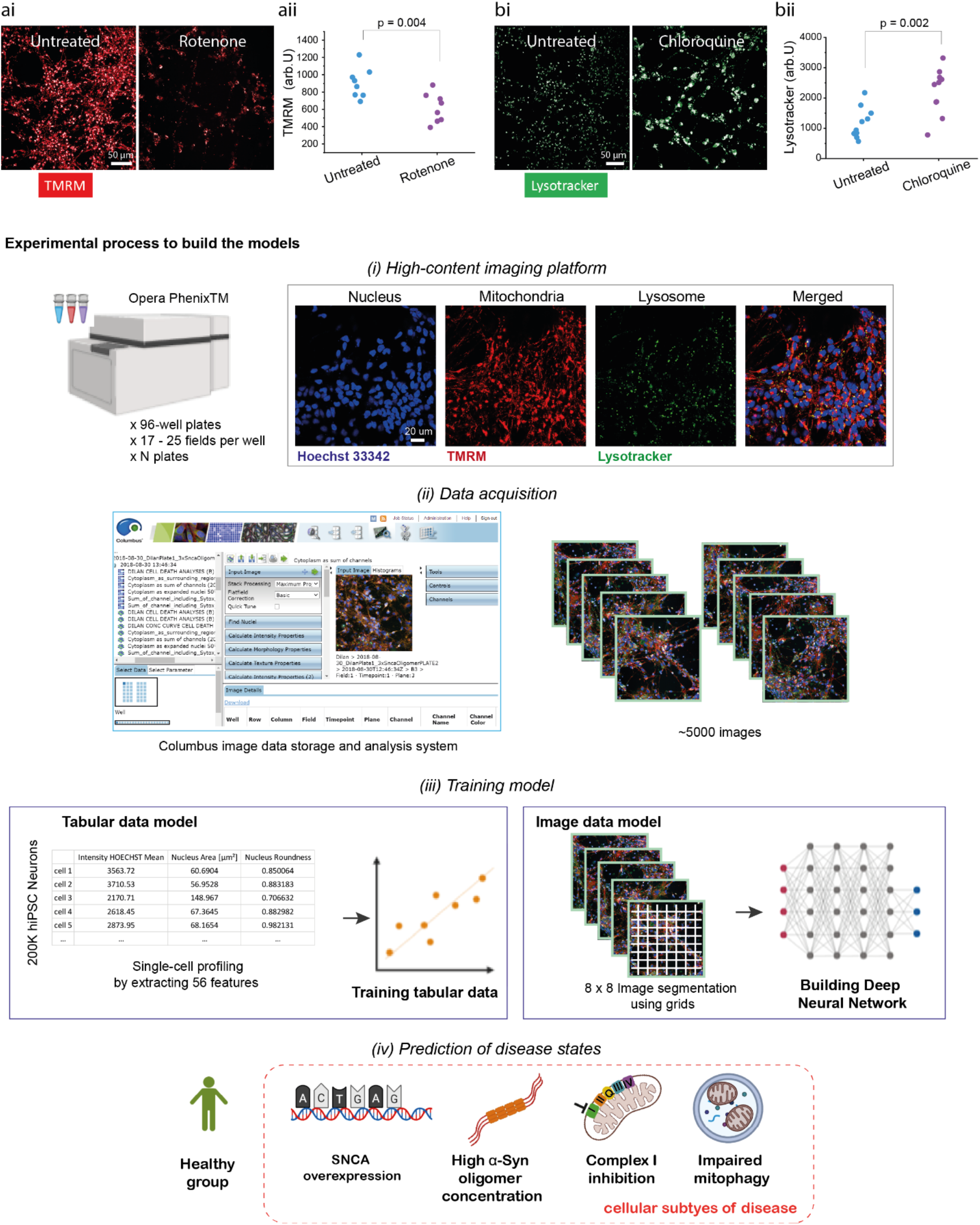
Workflow to develop a classifier to make a prediction of cellular subtypes in PD. **a & b.** High-throughput imaging enables visualization of mitochondrial depolarization by Complex 1 inhibitor, Rotenone (5 µM, ai & ii) and lysosomal activation by Chloroquine (1 µM, bi & ii). **c-f**. A schematic illustration to describe the experimental process to build the models. **c**. *(1)* Live-cell imaging with Opera Phenix high-content screening system (PerkinElmer): cells are loaded with live-cell imaging dyes. Representative images for the three channels; Hoechst 33342; nucleic labelling within 387/11 nm excitation and 417-447 nm emission, TMRM; mitochondrial labelling within 505 nm excitation and 515 nm emission, Lysotracker deep red; lysosomal labelling within 614 nm excitation and 647 nm emission. *(2)* Columbus Image Data Storage and Analysis System (PerkinElmer) was used to extract 56 morphological features (SFigure 1a & 2a) and whole images. *(3)* Models are trained on tabular data extracted from cell profiling features or images uniformly gridded by 8 × 8 segmented cropped images and categorically labelled and fed into the neural network. *(4)* The learned model enables the prediction of healthy or four disease subtypes (SNCA x3 genes, α-Syn Oligomer, mitochondrial Complex 1 inhibition and induction of Mitophagy based on primary PD pathology as described in Table 1.

To collect the dataset, the hiPSC derived neurons were labelled with live-cell imaging dyes, Hoechst3334, TMRM and Lysotracker deep red to label nucleus, mitochondria and lysosome respectively. Cell images were obtained using a high-content screening system to generate a large high-quality dataset. We also used SytoxGreen (labeling cells that have lost membrane integrity indicating cell death) to define a ‘live cell’ that describes the three organelle features in the same single cell unit by setting a fluorescent threshold (A.U 500 < live cells). We collected the data for AI models across a range of neuronal inductions (n = 6) and different plates (n = 8) in order to capture the inherent variation between cells, the hiPSC system, and the variation in conditions for dye loading and cellular imaging. We acquired the data in two formats 1) tabular data, which consisted of 56 mitochondrial, lysosomal, and nuclear features extracted from the Columbus Image Data Storage and Analysis System (PerkinElmer) and 2) 1024 × 1024 raw images, which we further segmented into smaller 8 × 8 tiles. We generated tabular and image models to identify disease status and the cellular subtypes of PD in human cells. These models consist of five classes; one control and four disease subtypes in an ‘all-in-one’ model (experimental process is described in Figure 2c).

### A classifier trained on cell profiles of key organelles predicts disease states

We built a classifier to predict five classes; four disease and one healthy state, using tabular data based on the nucleus, mitochondria and lysosome features extracted from the live high-content imaging platform. Organelle features extracted included cell areas, expression intensity, the number of spots, roundness, length and width. We also included SER textural features which measure local patterns of pixel intensity providing the structural information of the organelle loading (reviewed here (Di Cataldo and Ficarra, 2017) (Cretin et al., 2021). We designed, trained and evaluated a Dense Neural Network by splitting the entire dataset (n = 1,560,315 identified cells) into training (40%), validation (30%) and test datasets (30%). The confusion matrix demonstrates that overall, the correct label was identified 82% of the time in the five-class model. However, whilst some of the classes had an accuracy close to the model’s overall accuracy (SNCA x3: 84%, Oligomer: 82%, Mitophagy: 81%), specific disease subtypes had very high accuracy, notably complex 1 dysfunction (Complex 1: 98%), whilst the control state had slightly lower accuracy (Control: 69%, Figure 3bi). The accuracy of the model with all five classes was over 80% (SFigure 1bi), which was cross-validated using Stratified K-Fold Cross Validator (accuracy 81.3 ± 0.32 %, Figure 3bii), where the split of the dataset was randomized 10 times independently to validate the accuracy of the model. Next, we explored the feature importance of the model to understand the cellular features that drive accurate prediction using SHapley Additive exPlanation (SHAP) values on the entire test set. SHAP is an estimated value of importance for each feature in the model, and one dot represents each feature per cell (Lundberg et al., 2017). The ranked features based on their SHAP values, colored by their importance for each class, indicates that the majority of the features that explain the model’s predictions originate from mitochondria terms (‘TMRM’ related features, Figure 3c. Details of the features are described in SFigure 1a). This is highlighted for all disease subtypes where TMRM readouts are the highest SHAP based features, followed by the nucleus and lysosome features (Figure 3c). We then explored the top 10 features with the highest importance for each disease subtype individually to ascertain which organelle is driving the prediction for each pathway, and to understand the relationship between feature value and SHAP value (Figure 3di-v). Importantly, this demonstrates that out of the top 10 features for subtype 1 (SNCA x3), 5 were lysosomal, 3 were mitochondrial, and 2 were nuclear (Figure 3di); for subtype 2 (Oligomer), 7 were mitochondrial, 2 were nuclear, and 1 was lysosomal (Figure 3dii); for subtype 3 (Complex 1), 7 were mitochondrial, 2 were lysosomal, and 1 was nuclear (Figure 3diii); and for subtype 4 (Mitophagy), 6 were mitochondrial, were nuclear, and 2 were lysosomal (Figure 3div). Interestingly, this shows that whilst mitochondrial features are clearly critical to the model’s prediction, nuclear and lysosomal features are also important, with each featuring in the top 10 features for all disease subtypes. Finally, we selected the two most important features to the model’s overall prediction, two mitochondrial textural features (TMRM SER Valley and TMRM SER Dark), to assess if there is a significant difference across the five classes. The scatter graphs demonstrate that there is a significant difference in these features across the five classes (Figure 3ei & ii, *p* < 0.0001 between all groups), further demonstrating the importance of mitochondrial features in differentiating between disease states and healthy control.

**Figure 3.**
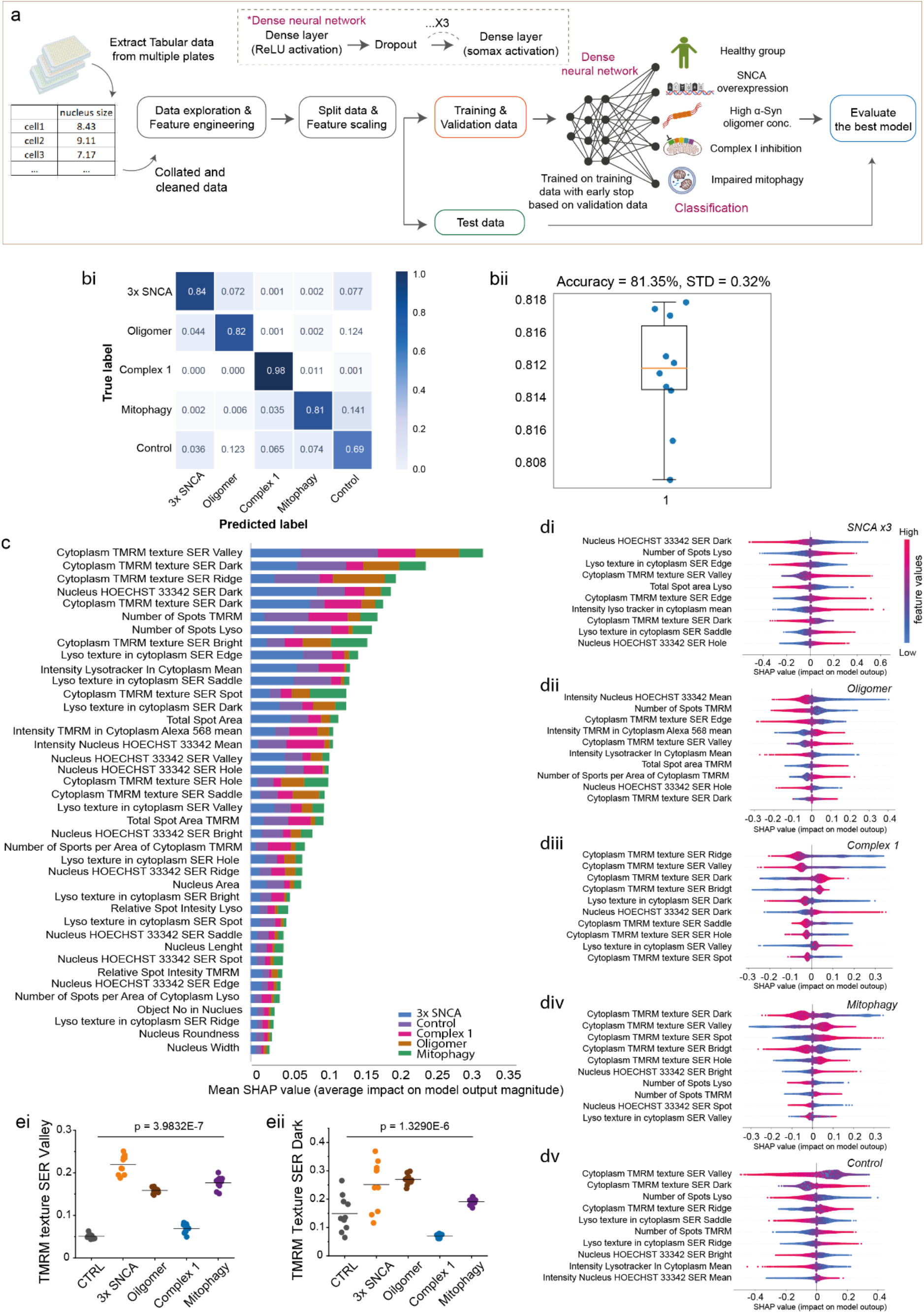
A classifier trained on cell profiles of key organelles predicts disease states with 82% accuracy. **a**. An illustration of workflow for machine learning with tabular data. **bi & ii**. Classification performance by a confusion matrix *(bi)* and the Stratified K-Fold Cross-Validation *(bii)* trained on cell profile tabular data. In the confusion matrix, the numbers from the confusion matrices denote the matching accuracy between true and predicted labels (true positive) and the diagonal elements correspond to precision. **c**. Feature ranking based on their SHAP values colored by class. **di-v**. A SHAP summary plot for the top 10 most important features based on their SHAP values for each of the classes: 3x SNCA (*di*), Oligomer (*dii*), Complex 1 (*diii*), Mitophagy (*dvi*) and Control (*dv*). Dots are colored according to the values of features for each cell, red represents high feature values, and blue represents lower feature values. A positive SHAP indicates an increased probability to predict each state (‘positive’ impact on the output) and vice versa. **e**. Random selection of 10 wells to test top two features measuring local patterns of pixel intensity shows an effect of cellular subtype across 5 groups (One-Way ANOVA *p* < 0.0005, n = number of wells). **Note**. Control: healthy group, 3x SNCA: SNCA mutation, Oligomer: treatment with α-Syn Oligomer, Complex 1: treatment with mitochondrial Complex 1 inhibition, Mitophagy: co-treatment with Antimycin and Oligomycin to induce Mitophagy.

### A model of the interaction between organelles predicts aggregation vs mitochondria phenotypes

In PD, the contact between mitochondria and lysosome is known to play a critical role in the homeostasis of organellar networks within the cell, for both damaged mitochondrial clearance (mitophagy) as well as lysosomal degradation of mitochondrial-derived vesicles. We demonstrate the presence of organellar contacts between mitochondria and lysosomes using super resolution microscopy in Figure 4a (Wong et al., 2018). Therefore, we reasoned that the contacts between mitochondria and lysosomes, captured in the multiplexed imaging platform, may reflect the index of mitophagy, and therefore provide an additional feature to predict mechanistic subtype. Of note, the use of a membrane potential dye such as TMRM will preferentially detect healthier mitochondria due to the nature of the dye loading.

**Figure 4.**
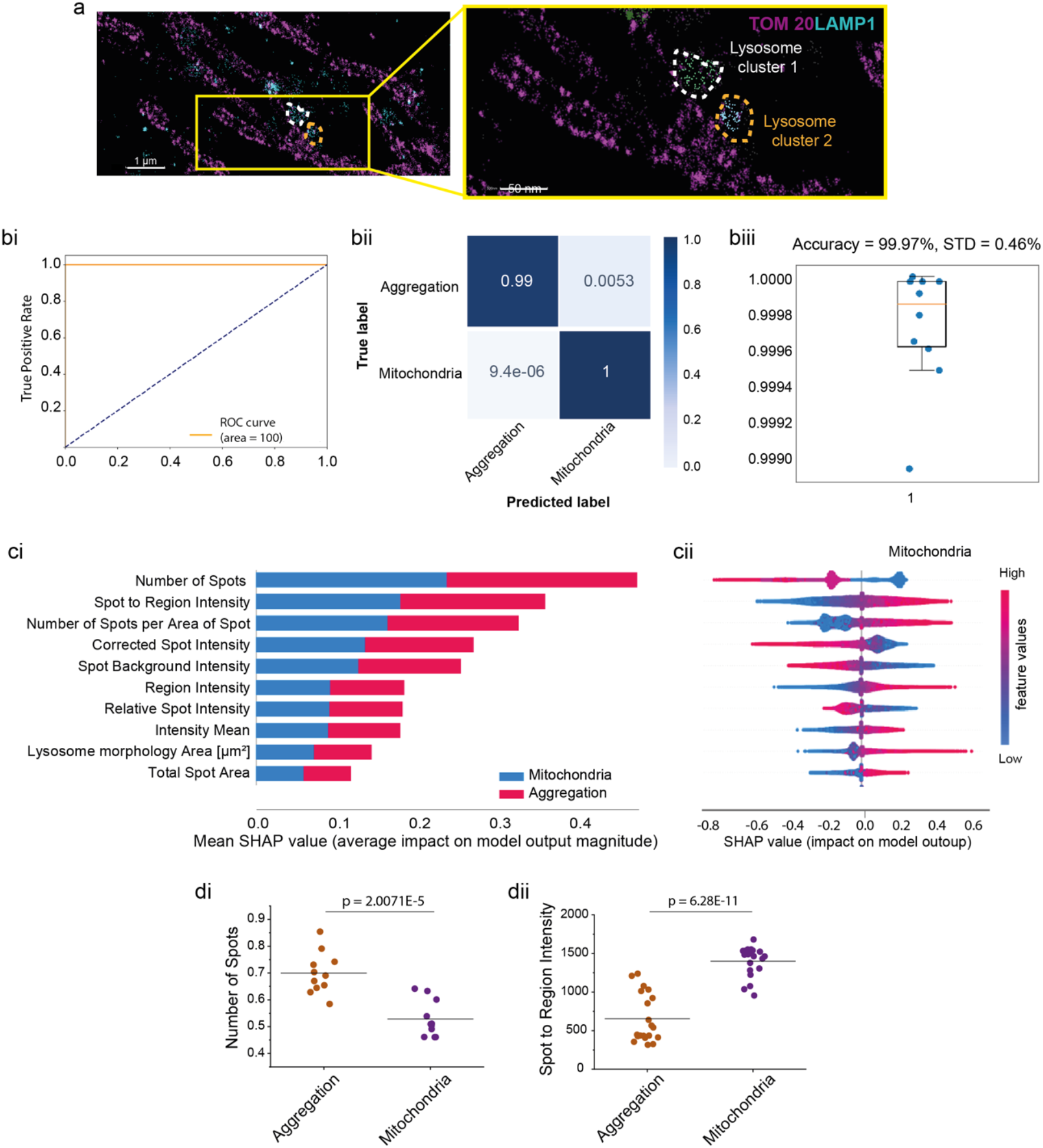
Interaction between cellular organelle networks classifies into aggregation and mitochondrial toxicity phenotypes. **a**. Super-resolution images of mitochondrial and lysosomal co-localization. Mitochondria are labeled with TOM 20 and lysosome with LAMP1. **bi-iii**. ROC-AUG curve of classification performance (*bi*), the confusion matrix (bii) and by Stratified K-Fold Cross Validation (*biii*) trained on the selected cell profile tabular data (mitochondria and lysosome contact). The selected tabular data from mitochondria and lysosome colocalization showing highly accurate prediction to identify the two disease states of mitochondrial toxicity and aggregation (> 99% accuracy). In the confusion matrix, the numbers from the confusion matrices denote the matching accuracy between true and predicted labels and the diagonal elements correspond to precision. **ci**. Feature ranking that drives the prediction of aggregation based on their SHAP values colored by their importance. **cii**. A SHAP summary plot of top 10 features to classify the groups into mitochondrial toxicity (aggregation group is shown as a mirror of this SHAP values, and therefore it is not presented). **di & ii**. Random selection of 12-22 wells to compare the top two features (‘Number of Spots’ indicates the number of lysosomal spots within mitochondria, and ‘Spot of Region Intensity’ indicates the lysosomal intensity co-localized with mitochondria) showing that there is a statistical significance between mitochondrial toxicity and aggregation groups (Two Sample test *p* < 0.0001). **Note**. Aggregation: combining subtypes of 3x SNCA and Oligomer. Mitochondrial toxicity: combining subtypes of Complex 1 and Mitophagy.

Here we trained a ML model using overlapping mitochondrial and lysosomal features, that is, the lysosomal spots that are within mitochondrial areas. We tested whether the model can distinguish the two primary pathologies of aggregation and mitochondrial dysfunction: we combined the SNCAx3 mutation and oligomer treatment into one single aggregation class, and combined the mitochondrial complex I function and mitophagy into one single mitochondria class. (aggregation vs mitochondrial toxicity; cell profile features are shown in SFigure 2a). This model predicted the correct mitochondrial label with 100% accuracy, and 99% accuracy for aggregation labels highlighting that the organellar interactions are highly informative in differentiating mitochondrial toxicity from aggregation toxicity in PD (Figure 4bi &ii). We cross-validated this result using Stratified K-Fold Cross Validator, observing an accuracy of 99.97 ± 0.46% across 10 independent runs (Figure 4biii). The model based on the interaction of organelles alone, showed a much lower performance in classifying the subtypes of the two main pathways (see the confusion matrix, SFigure 2d). We next looked at this feature importance between the two primary pathologies in more detail. SHAP analysis showed that the number of lysosomal spots and the lysosomal spot to region intensity within the mitochondria are the key features driving the accurate prediction between aggregation and mitochondrial toxicity (Figure 4ci & ii). This suggests that lysosomal contacts with the mitochondria may be a key feature in both aggregation and mitochondrial forms of PD. Finally, by plotting the top two features identified across plates, we show that there is an also significant change between those two forms of PD. Together, we show that a model based on the interaction between two organelles may be a predictor of broad disease type, and that data from both organelles can be used to accurately distinguish and predict aggregation from mitochondrial pathologies in PD.

### Convolutional Neural Network accurately predicts the disease states

We then created an image based classifier to distinguish between the different disease subtypes. The images were split into 5 classes; four disease pathways and one control as previously done for the tabular data-based classifier. The images were sliced into 8 × 8 tiled images, containing 1-20 cells per image, allowing preservation of the information contained in neuronal projections and the cell-cell contacts immediately surrounding neurons (reviewed here (Petralia et al., 2015)). We then trained a convolutional neural network to distinguish the 5 classes (see the schematic shown in Figure 5a). The confusion matrix from this model shows a true positive rate of > 90% for all five classes (average accuracy 95%, 3x SNCA: 89%, Oligomer: 99%, Complex I: 99%, Mitophagy: 94%, Control: 94%, Figure 5bi), highlighting a high accuracy to classify the disease states and control, especially when compared to the model trained on cell profiling tabular data (average accuracy: 82%). We cross-validated our results using Stratified K-Fold Cross Validator and showed 95.7% accuracy (Figure 5bii).

**Figure 5.**
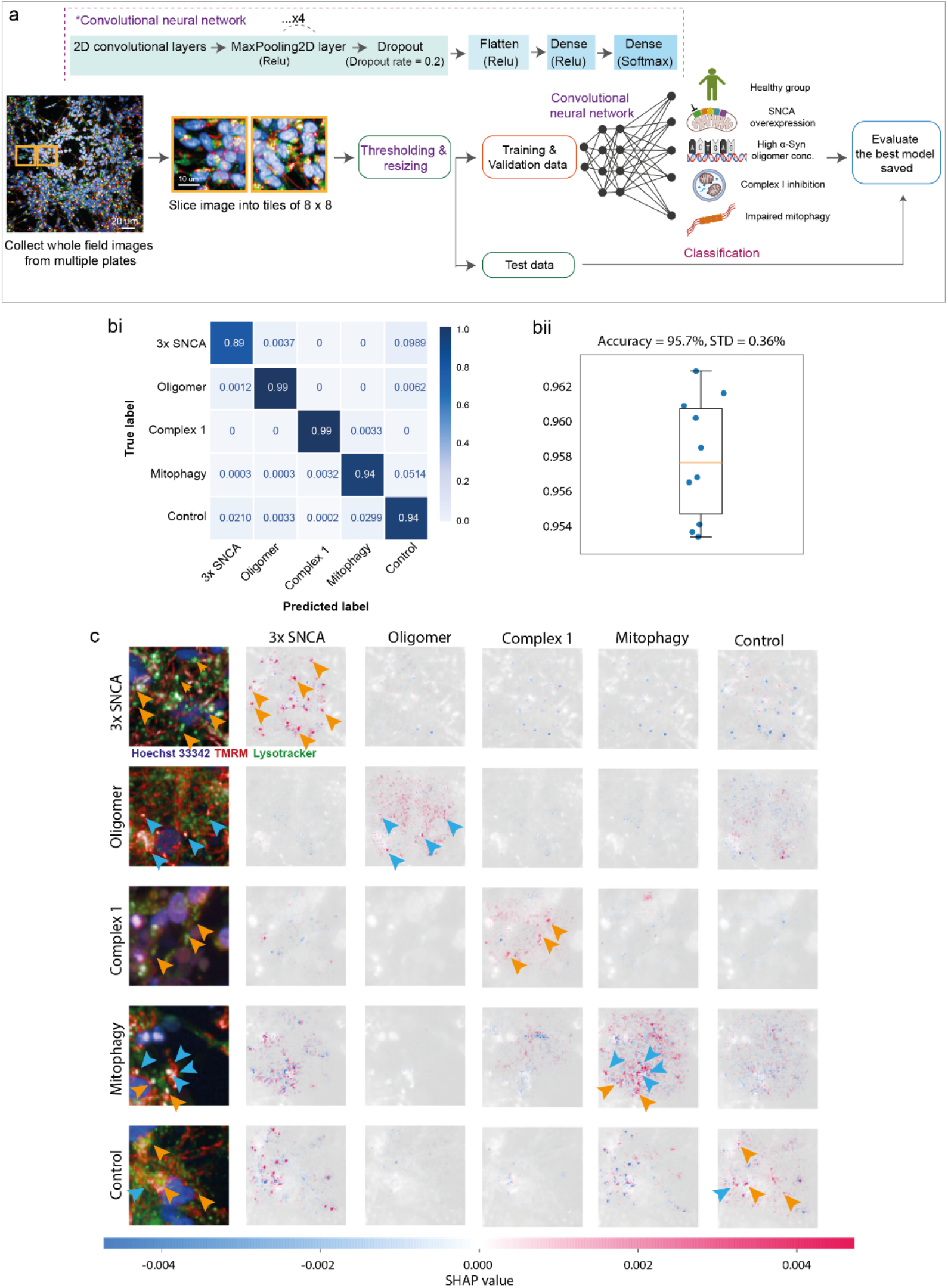
A classifier trained by images using Deep Neural Network accurately discriminates PD pathology. **a**. Illustration of workflow for Deep learning with images. **bi & ii**. Deep learning classification performance trained on 8 × 8 tiled images by the confusion matrix (*bi*) and the Stratified K-Fold Cross-Validation (*bi*). A sample is assigned to five classes with the maximum prediction accuracy (95%). **c**. A SHAP DeepExplainer plot summary. Rows show images from the test set, one from each class, and the columns represent each class. The SHAP value for each score is shown below. Orange and blue arrows indicate either lysotracker (lysosome) or TMRM (mitochondria) positive areas respectively. **Note**. Control: healthy group, 3x SNCA: SNCA mutation, Oligomer: treatment with α-Syn Oligomer, Complex 1: treatment with mitochondrial Complex 1 inhibition, Mitophagy: co-treatment with Antimycin and Oligomycin to induce Mitophagy.

We next employed the SHAP DeepExplainer method, which indicates how much each pixel contributes to the probability positively or negatively, in an attempt to understand which aspects of the image the model is using to make its predictions (Lundburg et al., 2007). It confirms that mitochondria along with the lysosomes are the major contributors driving the accurate prediction, and not the nucleus nor the negative space between cells (Figure 5c). Given the consistent result of the essential role of mitochondria to classify the disease states, we lastly explored whether images of mitochondria alone could accurately predict the disease states. We trained on images of mitochondria or lysosomes alone or the combination of both organelles. The true positive rate of models created for mitochondria is 87% (Figure 6ai, SFigure 4a), 82% for only lysosomes (Figure 6bi, SFigure 4b) and 90% for the duet images (Figure 6ci, SFigure 4c) and it was cross-validated using Stratified K-Fold showing above 82% accuracy (Figure 6aii, bii and cii). Taken together, we show that a deep neural network using image data provides a much more robust way to differentiate subtypes of PD, and identify which organelle is driving the pathology.

**Figure 6.**
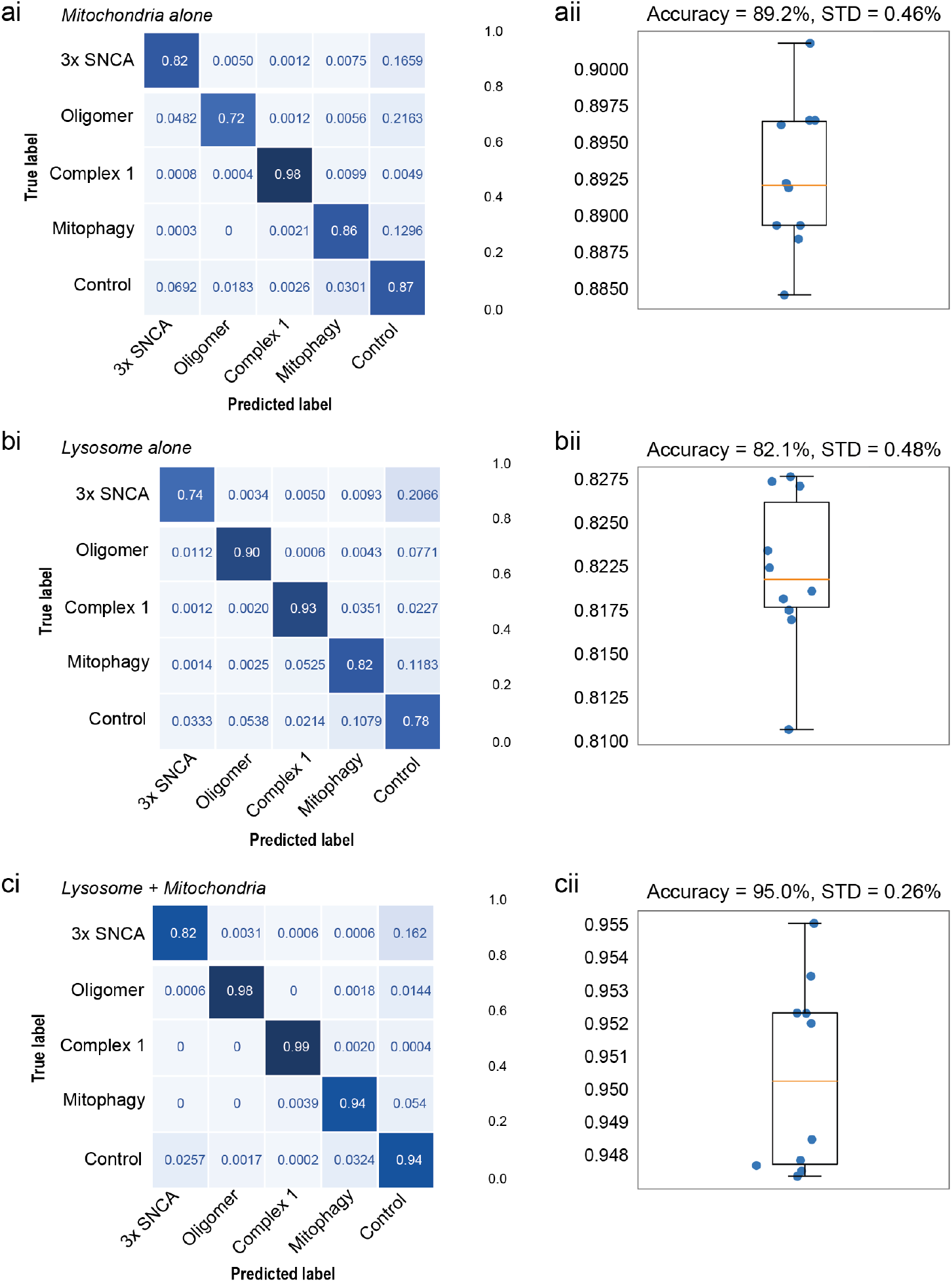
Deep Neural Network using mitochondria image alone retains high prediction accuracy. **a-c**. Deep learning classification performance the confusion matrices on trained on 8 × 8 tiled images and the Stratified K-Fold Cross-Validation of mitochondria alone (*ai & ii*), lysosome alone (*bi & ii*) and both together *(ci & ii*). **Note**. Control: healthy group, 3x SNCA: SNCA mutation, Oligomer: treatment with α-Syn Oligomer, Complex 1: treatment with mitochondrial Complex 1 inhibition, Mitophagy: co-treatment with Antimycin and Oligomycin to induce Mitophagy.

**Figure 7.**
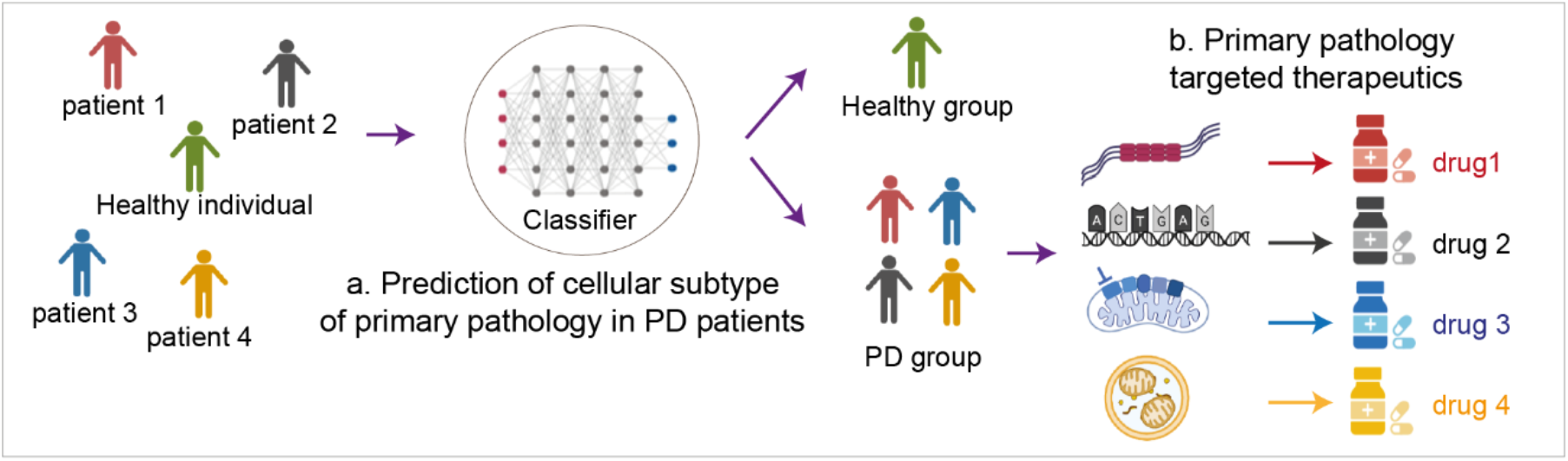
A schematic illustration demonstrates how machine learning based classifiers can be applied to improve the approach to PD therapeutics. **a**. Our classifier can classify individuals into PD and healthy groups. The PD diagnosed individuals can be further classified based on their mechanistic subtype. **b**. Mechanism-specific targeting drugs can be matched with the PD patients based on their own disease subtypes.

## DISCUSSION

Genome-wide association studies have identified multiple genetic risk loci relating to protein homeostasis, protein trafficking, lysosomal function and mitochondrial function in sporadic disease, implicating these pathways in disease pathogenesis (Chang et al., 2017; Ivatt et al., 2014; Nalls et al., 2011). Here, we used human iPSC-derived cortical neurons, a vulnerable cell type in PD, that robustly recapitulate critical cellular phenotypes of PD to successfully model and define four mechanistic subtypes of disease based on the presence of a familial mutation, proteotoxic stress, mitochondrial stress, and induced mitochondrial clearance. Using live high-content imaging, we tracked these disease mechanisms through three key organelles. Our approach is well placed as a preclinical platform to have high predictive value for disease as it is a human model of brain disease in a dish which captures live information on the two critical organelles implicated in PD. This convergence of benefits in our approach enabled us to then develop a highly accurate deep learning classifier, able to distinguish the presence or absence of disease, and if diseased, the subtype of the disease.

Using a range of intensity, morphological, and textural features of the mitochondria, lysosomes, and nuclei, extracted from the live high-content imaging platform, we trained a model to identify the primary pathology in hiPSC-derived cortical neuronal cultures. We demonstrate that this method is able to accurately predict between control cells and the four disease pathways. Importantly, the use of tabular data in this study enables the ranking of organellar features based on their contribution to the model’s prediction (using SHAP), providing unique and critical insight into the importance of mitochondrial and lysosomal biology in disease. This explainability demonstrates that mitochondrial features contribute most prominently to the overall prediction of the classifier, and specifically the prediction of mitochondrial pathways (Complex 1 and Mitophagy), with lysosomal and nuclear features also important in the prediction of aggregation pathways (3x SNCA and oligomer). Furthermore, when using a similar approach for the interaction between mitochondria and lysosomes we demonstrate that tabular data solely on the contacts between these organelles is sufficient to distinguish the two key categories: aggregation and mitochondrial toxicity pathways with high accuracy. Taken together, these classifiers demonstrate that the information contained within the mitochondria and lysosomes independently, and the information contained within their interaction (for example when lysosome clear mitochondria), are both sufficient to predict, and are therefore likely to be biologically relevant, in the four mechanisms in disease described here.

Whilst models trained on tabular data are advantageous due to the level of explainability they provide, they may be susceptible to minor alterations in experimental conditions, such as the loading of imaging dyes and the pre-processing of image data. They may also be dependent on the software processing that converts the images into tabular data which is subject to another filtering level that carries uncertainty with it. As a result, they may lack generalizability, and perform less well. We, therefore, also generated CNN-based image classifiers using the same large dataset for the tabular data-based model (Christiansen et al., 2018). We show that deep CNN-based image classifiers can correctly classify images to accurately identify a disease state from a healthy control cell, and this is more generalizable, and shows high performance achieving close to 100% accuracy. Nevertheless, both our tabular and image classifier approaches were able to accurately discern which underlying pathogenic pathways was driving the disease state

Our approach offers several advantages to current perspectives to analyzing cell images. Although traditional methods are capable of quantifying well-defined structural properties, and focus on automation and throughput, they do not capture all of the information contained within imaging data. Using traditional image processing software, researchers must typically first choose which feature (or features) to combine to quantify from a vast array of possible cellular phenotypes, such as nuclear fragmentation, protein aggregation, cell density, organelle damage and distribution, and changes to cytoskeleton organization. This approach can be challenging and time-consuming and may be subject to bias. An alternative approach described here uses machine learning to decipher cellular features at high accuracies in an unbiased manner and achieve a greater accuracy than using traditional throughput readouts. Similar use of CNNs have been used successfully to accurately classify images of iPSC spinal motor neurons taken from healthy controls and patients with amyotrophic lateral sclerosis (ALS) (Imamura et al., 2021).

The high accuracy achieved by our tool was made possible by several conditions. The cell density was relatively high (and necessary) in order to provide enough information for the training and validation sets, and also, we made use of image preprocessing to increase the number of images available to the CNNs. In addition, we trained CNNs, with multiple hidden layers to improve the accuracy. Despite the underlying complexity of the deep learning algorithms used, the methods and tool used to classify the images was relatively simple (Grafton et al., 2021). Only minimal image processing was needed, with each image being divided into 64 individual “tiles”. While training a CNN has been made simpler in recent years with the development of user-friendly software applications such as Tensorflow. In addition, the use availability and access to a GPU allowed the processing of large amounts of data in a relatively short time (somewhat necessary given the number of images to be processed).

In this study, our approach of creating multiple classifiers based on extracted tabular features and image tiles enables us to gain explainability from the tabular data and translate this to the image classifier by training models on mitochondria and lysosomes individually, and in combination. We demonstrate that the loss of nuclear signal, leaving both the mitochondria and lysosomes, in a CNN-image based model does not reduce the accuracy of the model. We further demonstrate that a CNN trained on mitochondria alone achieves higher accuracy than the model based on lysosomes alone, consistent with the feature importance generated from the tabular models.

Nevertheless, the approach outlined here demonstrates the power of using deep learning in predicting the underlying mechanisms of PD. Importantly, as PD is highly heterogeneous this platform may enable the disease mechanism in patient cells to be classified. This may have significant clinical implications in both diagnosis and treatment, as the identification of cellular mechanisms may indicate their likely response to a proteinopathy treatment (e.g. targeting α-synuclein) versus a mitochondrial treatment (e.g. antioxidant therapy). Furthermore, this approach establishes a drug discovery platform that may be used to assess which pathway predominates in an individual, and whether specific medications are capable of reversing robust cellular phenotypes, in an unbiased approach (Chandrasekaran et al., 2021).

## METHODS

### Generation of human iPSC

We generated neurons derived from human iPSCs (hiPSCs) that are reprogrammed from two healthy donors (one generated in house, and one commercial line (ThermoFisherScientific)) or PD patients carrying mutations (SNCA triplication, 3x SNCA) who had given signed informed consent for the derivation of hiPSC lines from skin biopsy as part of the EU IMI-funded program Stem-BANCC (Chen et al., 2019; Devine et al., 2011). All experimental protocols had approval from the London-Hampstead Research Ethics Committee and R & D approval from University College London Great Ormond Street Institute of Child Health and Great Ormond Street Hospital Joint Research Office. The hiPSCs were cultured on Geltrex (ThermoFisherScientific) in E8 media (ThermoFisherScientific) or mTeSR (Stem Cell Technologies) and passed using 0.5 mM EDTA (ThermoFisherScientific). All lines were mycoplasma tested (all negative) and short tandem repeat profiled (all matched) performed by the Francis Crick Institute.

### Generation of high content imaging data and image processing

We collected sets of z-stack images from various focal planes in the z-axis while the stage was fixed in the × and y-axis. To reduce undesirable effects of out-of-focus features, the training samples consist of two-dimensional information from maximum intensity projections of the z-stack images with fluorescence labels that are pixel registered. The cell profiles based on tabular data and the image dataset were separately collected using the Columbus image storage and analysis system.

### Classifiers trained

Tabular data was extracted from 7 plates consisting of four disease and control groups (60 % of control is labelled as healthy per plate) using the Columbus Data analysis and Storage system. For data exploration and feature engineering, the data points where half the features were missing were excluded. This was followed by splitting the dataset into training (n = 13489), validation (n = 28373) and test (n = 35466) datasets. The features (lists in SFigure 1a) were scaled per control in each plate separately using the Power Transformer scaler. The scaling factor, lambda, was examined in the training dataset, and features with high variance (lambda >100) were excluded (all features showed lower variances and, therefore, all included). We designed and trained a Dense Neural Network with python using Tensorflow (Model structure: 3 dense layers using ReLU activation, each followed by a Dropout layer with dropout rate of 0.2 and a final dense layer with softmax activation). We used Adaptive Moment Estimation as the optimizer, monitoring the validation loss to save the best model and plotting the training and validation losses and accuracy. We used SHAP values to explore the feature importance of our 5-class model with a view of gaining insight into the cellular features that drive accurate prediction.

The accuracy of the model was cross-validated using Stratified K-Fold Cross Validator, where the split of the dataset was randomized 10 times independently to validate the accuracy of the model. Briefly, data were split into 10 stratified folds preserving the percentage of samples of each class across the folds. The model was run 10 times with each fold being used as the test set and the remianing 9 folds being used as the training set. Each training set returned by StratifiedKFold was split further into training and validation datasets using the stratify parameter. The training, validation and test dataset went through the steps of feature scaling followed by creating and training the model as described in the single runs. The validation loss was monitored, and the best model was saved and evaluated on the test dataset. We then average the performance of the 10 test datasets and report the mean and standard deviation.

The data to highlight the interactions between mitochondria and lysosomes, was generated with single-molecule localization microscopy (resolution of 20 nm) to visualize the contacts. The tabular data (lists in SFigure 2a) generated from 5 plates (number of cells; controls = 845,143 mitophagy = 644,457, 3xSNCA = 101,158, a-Syn oligomer =33,671, Complex 1 = 15,751) were processed for data exploration and feature engineering. We then split the data into training (n = 1,049,715), validation (n = 328,036) and test (n= 262,429) datasets. The features were scaled per control in each plate separately using the Power Transformer scaler. The scaling factor, lambda, was examined in the training dataset and one feature (Lysosome texture SER Hole 0 px) with high variance across plates (lambda >50) was excluded from the training, validation and test datasets. The controls were excluded from the datasets after feature scaling resulting in n= 508,415 in the training dataset, 159,234 in the validation and 127,388 in the test dataset. We designed and trained a Dense Neural Network using Tensorflow, with the same model structure described for the tabular data above and illustrated in Figure 3a. We trained with a batch size of 256 and stopped if the validation loss did not improve for 50 consecutive epochs or the number of epochs exceeded 500. The same model structure was used to build the classifier to predict the 5 classes.

For image models, each of the 1024 × 1024 pixels high-throughput images, consisting of 100-400 neuronal cells (each neuron consists of approximately 900-22,500 pixels), was sliced into an 8 × 8 tiled image; 3x SNCA (n = 7983), Oligomer (n = 8307), Complex 1 (n = 11875), Mitophagy (n = 13692), and one control group (n= 22,461) that contained 1-20 cells per sliced image. After cropping, dark images due to few numbers of cells contained were removed by applying a cut-off of 0.1 on the mean intensity threshold and a variance threshold of 0.0275. The image tiles were shuffled and resized to 84 × 84 and split into training (n= 51,454) and test (n= 12,864) datasets. The training data was then further split into training (n = 41,163) and validation (n = 10,291) datasets and shuffled before batching. The neural network was implemented using Tensorflow (see Figure 5a for the architecture).

### Data and software availability

Image processing pipelines, all tabular data and whole images (before tiling) are publicly available as deposited in Zenodo under the access number 5823423 (DOI: 10.5281/zenodo.5823423). The codes for the models are available at https://github.com/Minee-Liane-Choi/Gandhi-PD-Classifier-2022.

## ACKNOWLEDGEMENTS

SG is an MRC Senior Clinical Research Fellow (MR/T008199/1). This work was also supported by the funding from the Francis Crick Institute ITO (Information Technology Office). We would like to thank Daniella Melandri from UCL Queen Square Institute of Neurology and Dr Michael Howell and Dr Ok-Ryul Song from the High-Throughput Screening team of the Francis Crick Institute for their advice in building analysis pipelines.

## AUTHOR CONTRIBUTIONS

Conceptualization, M.L.C., S.G.; Data collection and analysis, D.A., M.L.C.; Investigation, K.D., J.R.E., G.S.V.; Technical support., G.V., A.A., O.B., J.F., Writing original manuscript, K.D., J.R.E., G.S.V., M.L.C, D.A.; Writing review & editing., S.G., M.L.C., D.A., G.V., A.A., O.B., H.C.; Funding Acquisition, S.G.

## SUPPLEMENTARY FIGURES

**SFigure 1.**
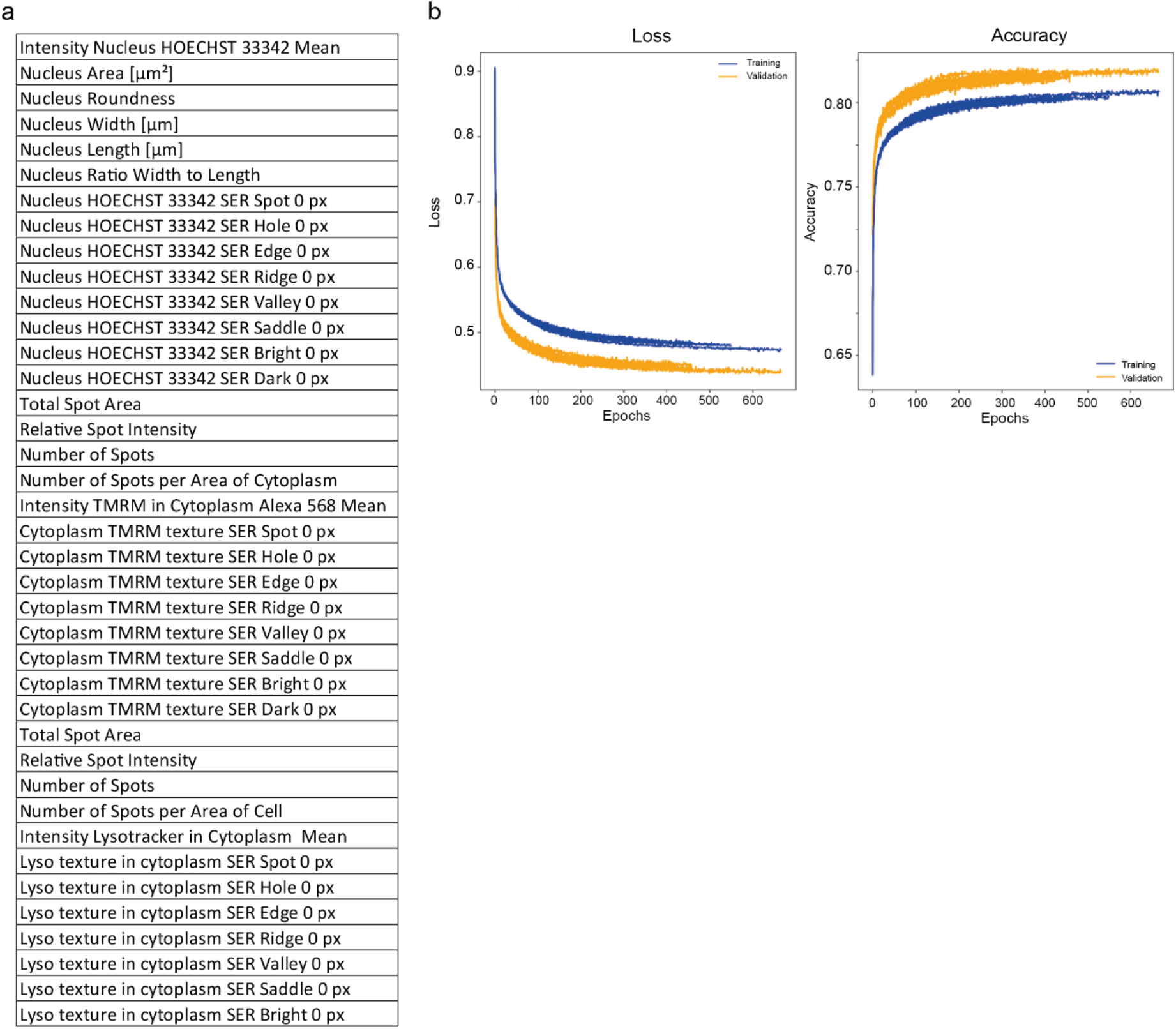
Supplementary data for the main Figure 3. **a**. The features regarding three key organelles, nucleus (Hoechst3337), mitochondria (TMRM) and lysosome (Lyso) used to train tabular data. Morphologically defined features are included such as cell areas, expression intensity, the number of spot, roundness, length and width. SER texture features are also included defined as Spot, Hole, Edge, Ridge, Valley, Saddle, Bright and Dark which measure local patterns of pixel intensity providing the structural information of the organelle loading (reviewed here (Di Cataldo and Ficarra, 2017) (Cretin et al., 2021). **b**. The Loss and Accuracy curve (training and validation).

**SFigure 2.**
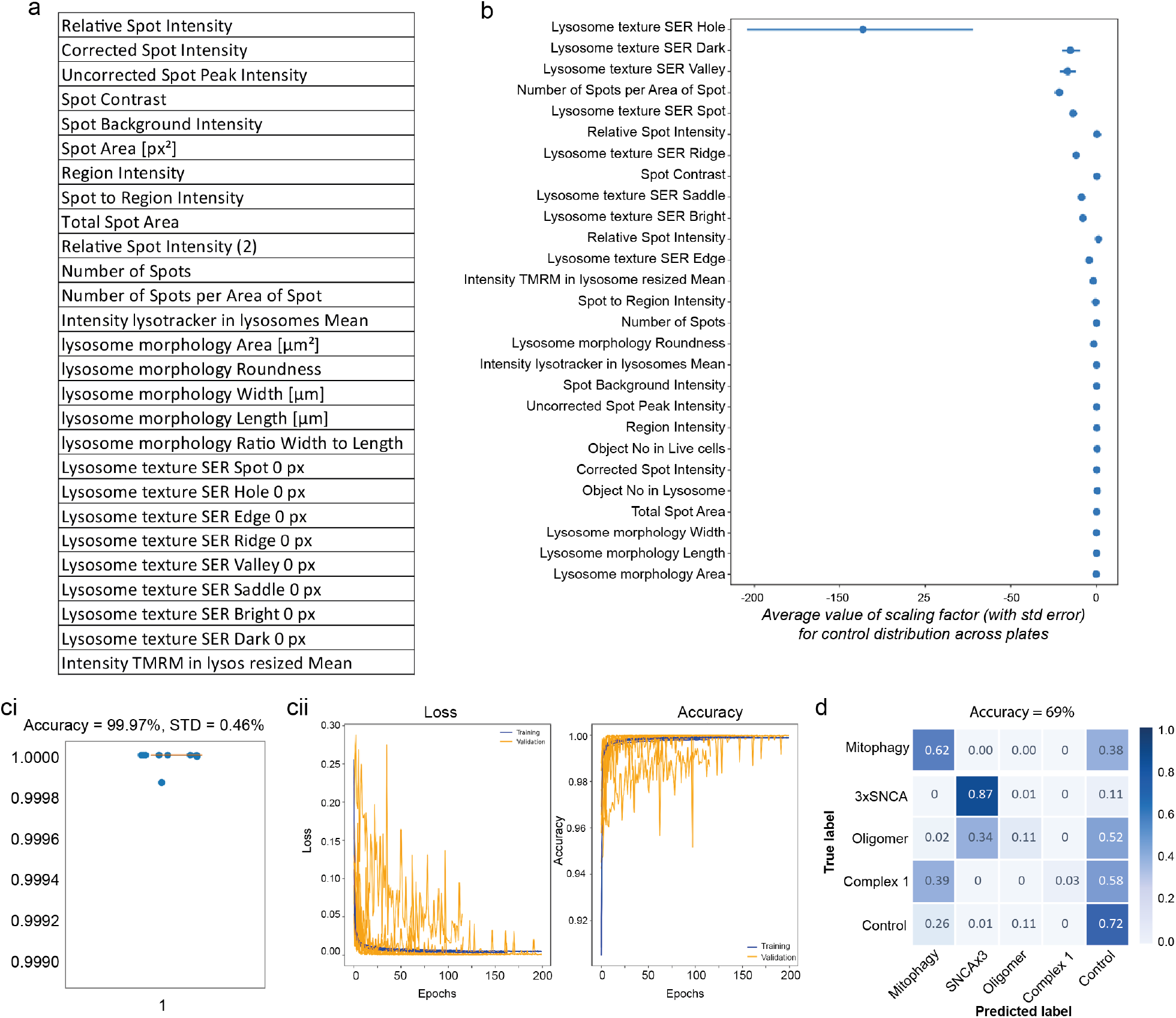
Supplementary data for the main Figure 4. **a**. Cell profiling features for the lysosomal features that contact mitochondria. **b**. The plot shows the average scaling factor, with standard error, for control distribution across plates for the lysosomal features that contact mitochondria in the training dataset. The features were scaled per control in each plate using the Power Transformer scaler. Those with a high variance in the scaling factor, lambda, (> 50) in the training dataset, such as ‘Lysosome texture SER Hole’ feature were excluded from the training, validation and test dataset. **ci & ii**. ROC-AUG (*ci*) and the Loss and Accuracy curve (training and validation, *cii*) by Stratified K-Fold Cross-Validation of the model to identify aggregation vs mitochondrial toxicity group. **d**. Confusion matrix of 5-classes model training on the selected data from the mitochondrial and lysosomal co-localization.

**SFigure 3.**
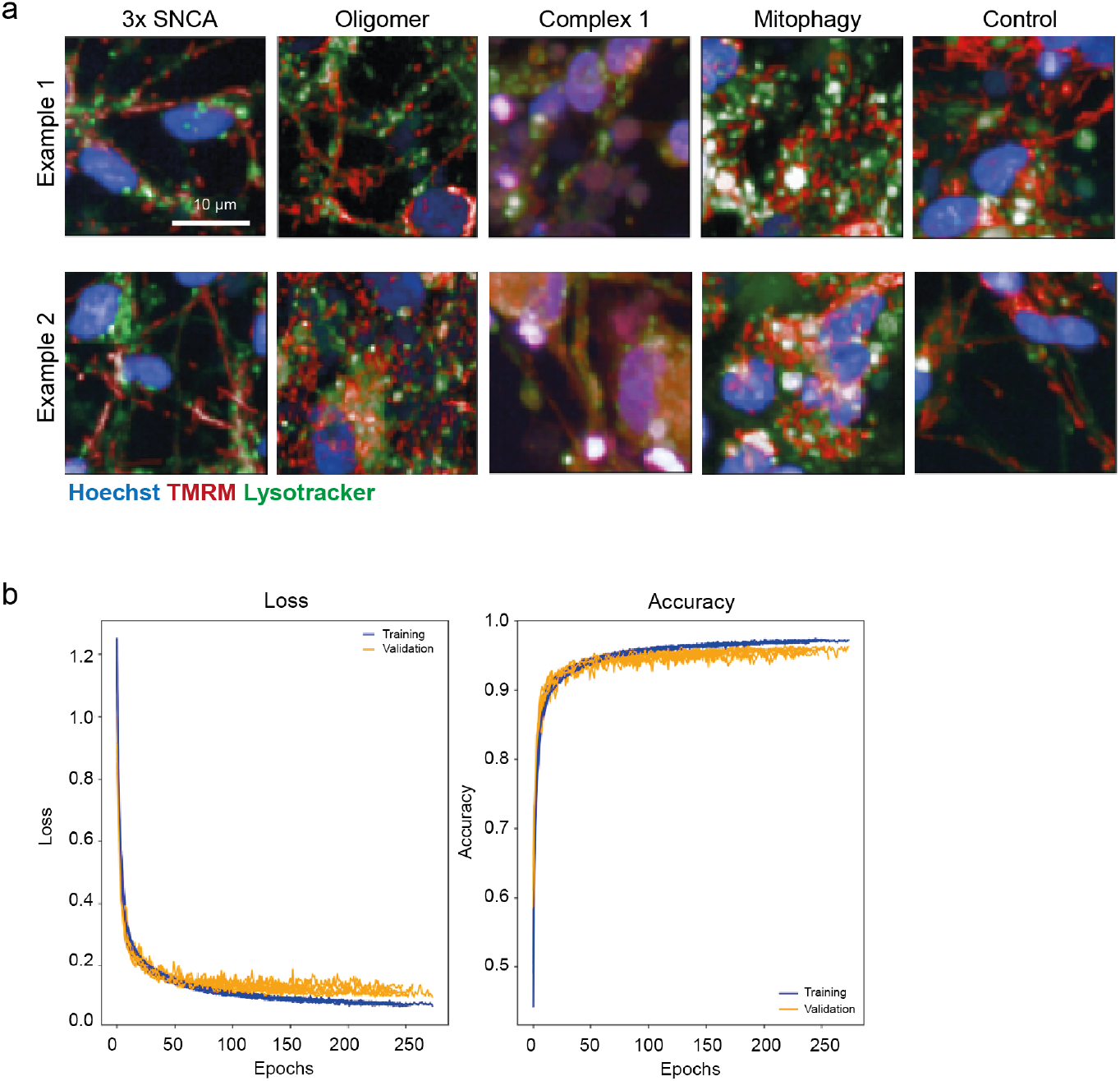
Supportive data for the main Figure 5. **a**. Examples of 8×8 sliced images that merged each of Hoechst, TMRM and Lysotracker images from the test set that are predicted above 99.99% accuracy from each class. **b**. The Loss and Accuracy curve (training and validation).

**SFigure 4.**
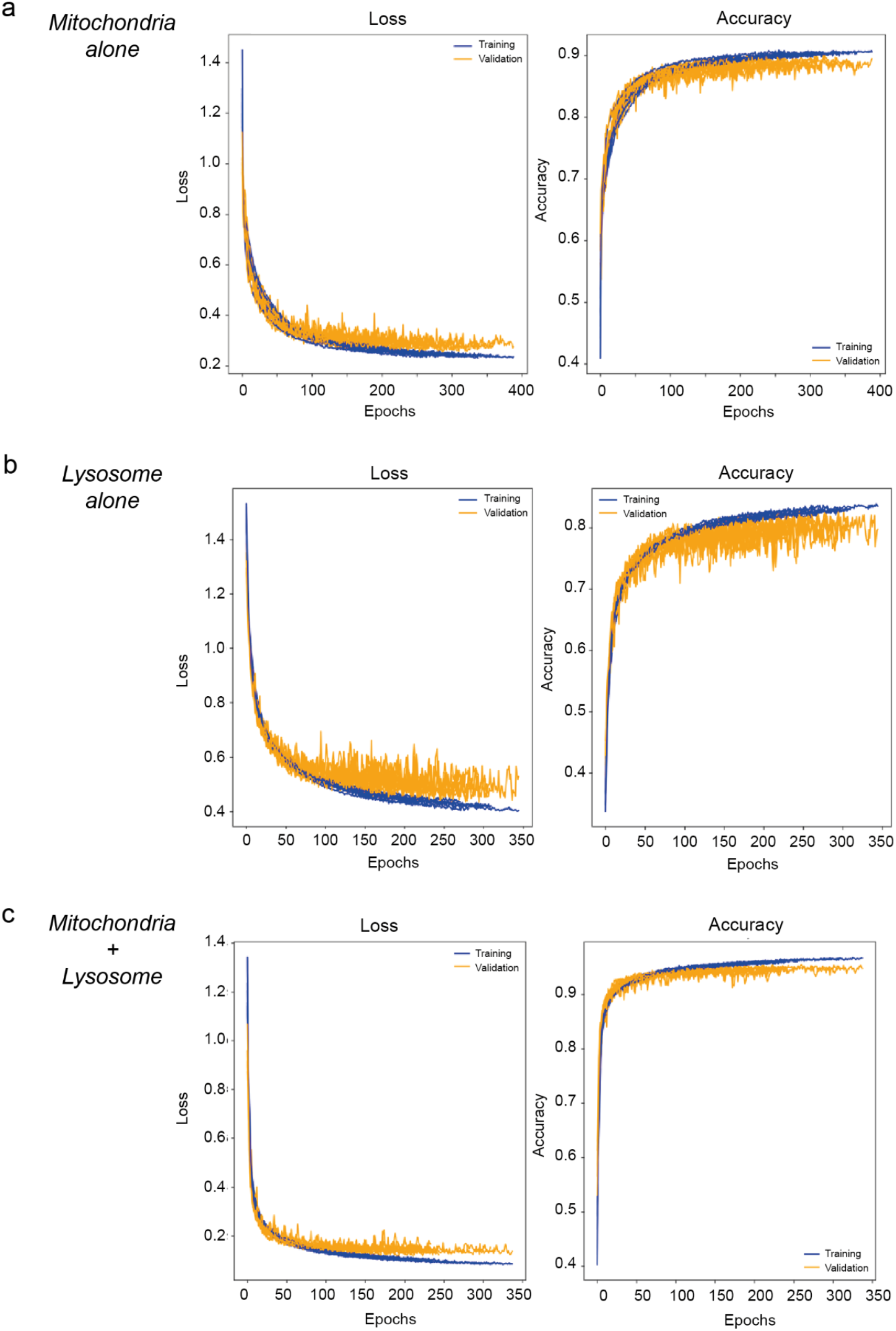
Supplementary data for the main Figure 6. The loss and accuracy curve (training and validation) of the tile-based images of mitochondria alone *(a)*, lysosome alone *(b)* and both together *(c)*.

